# The fitness cost of mis-splicing is the main determinant of alternative splicing patterns

**DOI:** 10.1101/114215

**Authors:** Baptiste Saudemont, Alexandra Popa, Joanna L. Parmley, Vincent Rocher, Corinne Blugeon, Anamaria Necsulea, Eric Meyer, Laurent Duret

**Author notes:** these authors contributed equally to this work (Baptiste Saudemont, Alexandra Popa, Joanna L. Parmley).

## Abstract

Most eukaryotic genes are subject to alternative splicing (AS), which may contribute to the production of functional protein variants or to the regulation of gene expression, notably via nonsense-mediated mRNA decay (NMD). However, a fraction of splice variants might correspond to spurious transcripts, and the question of the relative proportion of splicing errors vs. functional splice variants remains highly debated. We propose here a test to quantify the fraction of AS events corresponding to errors. This test is based on the fact that the fitness cost of splicing errors increases with the number of introns in a gene and with expression level. We first analyzed the transcriptome of the intron-rich unicellular eukaryote *Paramecium tetraurelia.* We show that both in normal and in NMD-deficient cells, AS rates (intron retention, alternative splice site usage or cryptic intron splicing) strongly decrease with increasing expression level and with increasing number of introns. This relationship is observed both for AS events that are detectable by NMD or not, which invalidates the hypothesis of a possible link with the regulation of gene expression. Our results indicate that in genes with a median expression level, 92%-98% of observed splice variants correspond to errors. Interestingly, we observed the same patterns in human transcriptomes. These results are consistent with the mutation-selection-drift theory, which predicts that genes under weaker selective pressure should accumulate more maladaptive substitutions, and therefore should be more prone to errors of gene expression.

## INTRODUCTION

The maturation of a primary transcript by the spliceosome can lead to the production of diverse transcripts, via the use of different splice sites and/or intron retention. Alternative splicing (AS) is widespread in eukaryotes, and it has been postulated that it might considerably expand the functional repertoire of eukaryotic genomes (Graveley 2001; Nilsen and Graveley 2010; Blencowe 2006). Many case studies have shown that some AS events are functional, i.e. that they play a physiological role, beneficial for the fitness of the organism (for review see (Kelemen et al. 2013)). However, like any biological machinery, the spliceosome is not 100% accurate, and the splicing of primary transcripts occasionally leads to the production of spurious mRNAs. These erroneous transcripts represent a waste of resources and may lead to the production of toxic protein variants, and hence are expected to be deleterious for the fitness of organisms. Indeed, several quality control mechanisms exist in eukaryotic cells to mitigate the negative impact of erroneous transcripts (Graille and Séraphin 2012). In particular, the nonsense-mediated decay (NMD) machinery is able to recognize and degrade cytoplasmic transcripts containing premature termination codons (PTCs) (Popp and Maquat 2013). However, these quality-control processes themselves are not 100% efficient. Hence, any transcriptome necessarily includes a fraction of variants that correspond to splicing errors, and their frequency relative to functional AS events remains open for debate.

In a large majority of cases, splice variants contain PTCs (i.e. encode truncated proteins), and only a very small fraction (<0.6%) of annotated AS events lead to the production of a detectable amount of protein (Abascal et al. 2015). Moreover, AS events are generally not conserved across species (Barbosa-Morais et al. 2012; Merkin et al. 2012; Reyes et al. 2013), and alternative splice sites show no sign of selective constraint (Pickrell et al. 2010). The subset of AS events that are strongly tissue-specific and that preserve the reading frame is generally more conserved, which clearly suggests that this subset includes some functional events. However these cases represent only a small fraction of all AS events (Barbosa-Morais et al. 2012; Merkin et al. 2012; Reyes et al. 2013). These observations indicate that the majority of AS events are not involved in the production of functional protein variants. This led some authors to conclude that the vast majority of AS events correspond to splicing errors (Melamud and Moult 2009; Pickrell et al. 2010; Wang et al. 2014; Stepankiw et al. 2015) (we will hereafter refer to this hypothesis as the “noisy splicing” model).

However, this interpretation is contested by other authors who argue that AS might play an important role in the regulation of gene expression. Indeed, the maturation of primary transcripts into PTC-containing splice variants, which then get degraded by NMD, can be used as a way to regulate the amount of mRNA available for protein production (this posttranscriptional regulation pathway is termed AS-NMD, for AS coupled with NMD; for review, see (McGlincy and Smith 2008; Hamid and Makeyev 2014)). AS-NMD notably plays an important role in the regulation of genes involved in the splicing process itself, presumably to maintain the homeostasis of splicing factors via auto-regulatory loops (Lareau et al. 2007; Ni et al. 2007). Interestingly, although the regulation of splicing factors by AS-NMD is well conserved across animals, the AS events that trigger NMD in these genes often involve different splice sites (Lareau and Brenner 2015). The rapid evolution of AS events in mammals is therefore not necessarily in contradiction with the hypothesis that many of them play an important regulatory role. The comparison of transcriptomes in normal vs. NMD-deficient cells revealed that a large fraction of genes produce splice variants (in a broad sense, i.e. including cases of intron retention) that are targeted by NMD (Ramani et al. 2009; Weischenfeldt et al. 2012; Kalyna et al. 2012; Drechsel et al. 2013; Stepankiw et al. 2015). This pattern is widespread in eukaryotes and is not restricted to genes encoding splicing factors. Importantly, patterns of AS vary among tissues and during cell differentiation (Wong et al. 2013; Braunschweig et al. 2014; Edwards et al. 2016). This led several authors to propose that AS-NMD might play a critical role in broadly regulating expression of a large percentage of genes (Wong et al. 2013; Braunschweig et al. 2014; Ge and Porse 2014; Smith and Baker 2015; Wong et al. 2016; Edwards et al. 2016).

Beyond a few case studies that provided clear evidence of genes regulated by AS-NMD, we still lack a global picture of the relative prevalence of functional AS compared to splicing errors. We propose here a test to quantify the fraction of splice variants corresponding to errors, i.e. having a negative impact on the fitness of organisms. The basis of this test is that the strength of splice signals is expected to reflect a balance between selection (which favors alleles that are optimal for splicing efficiency) and mutation and random genetic drift (which can lead to the fixation of non-optimal alleles) (Bulmer 1991). This selection-mutation-drift equilibrium therefore predicts a higher splicing accuracy at introns where errors are more deleterious for the fitness of organisms. Hence, if AS events predominantly correspond to splicing errors, one should expect a negative correlation between the rate of AS events and their cost in terms of resource allocation (metabolic cost, mobilization of cellular machineries). The noisy splicing model therefore makes several specific predictions regarding the AS rate according to whether splice variants are detectable by NMD or not, and according to the expression level, length and number of introns of genes.

We first implemented this test in the ciliate *Paramecium tetraurelia.* The genome of this unicellular eukaryote contains a large number of introns (> 90,000 introns; 2.3 introns per gene on average). One major advantage of this organism is that its introns are very short (25.1 bp on average, with 99.9% of them between 20 and 35 bp; Fig. 1A), i.e. much shorter than RNAseq sequence reads, which greatly simplifies the detection and classification of AS events. In particular, cases of intron retention can be identified directly by detecting sequence reads spanning the entire intron and its flanking exon boundaries (Fig. 1C). Moreover, this organism already proved to be a good model to reveal important general features of splicing control in eukaryotes (Jaillon et al. 2008). Here we present a comprehensive characterization of AS in the transcriptomes of normal and NMD-deficient paramecia to test the AS-NMD and noisy splicing models. We then ran the same test using previously published human transcriptome data sets, and we quantified the fitness cost of mis-splicing in humans by analyzing polymorphism data. Our analyses reveal that the vast majority of splice variants correspond to errors.

**Figure 1:**
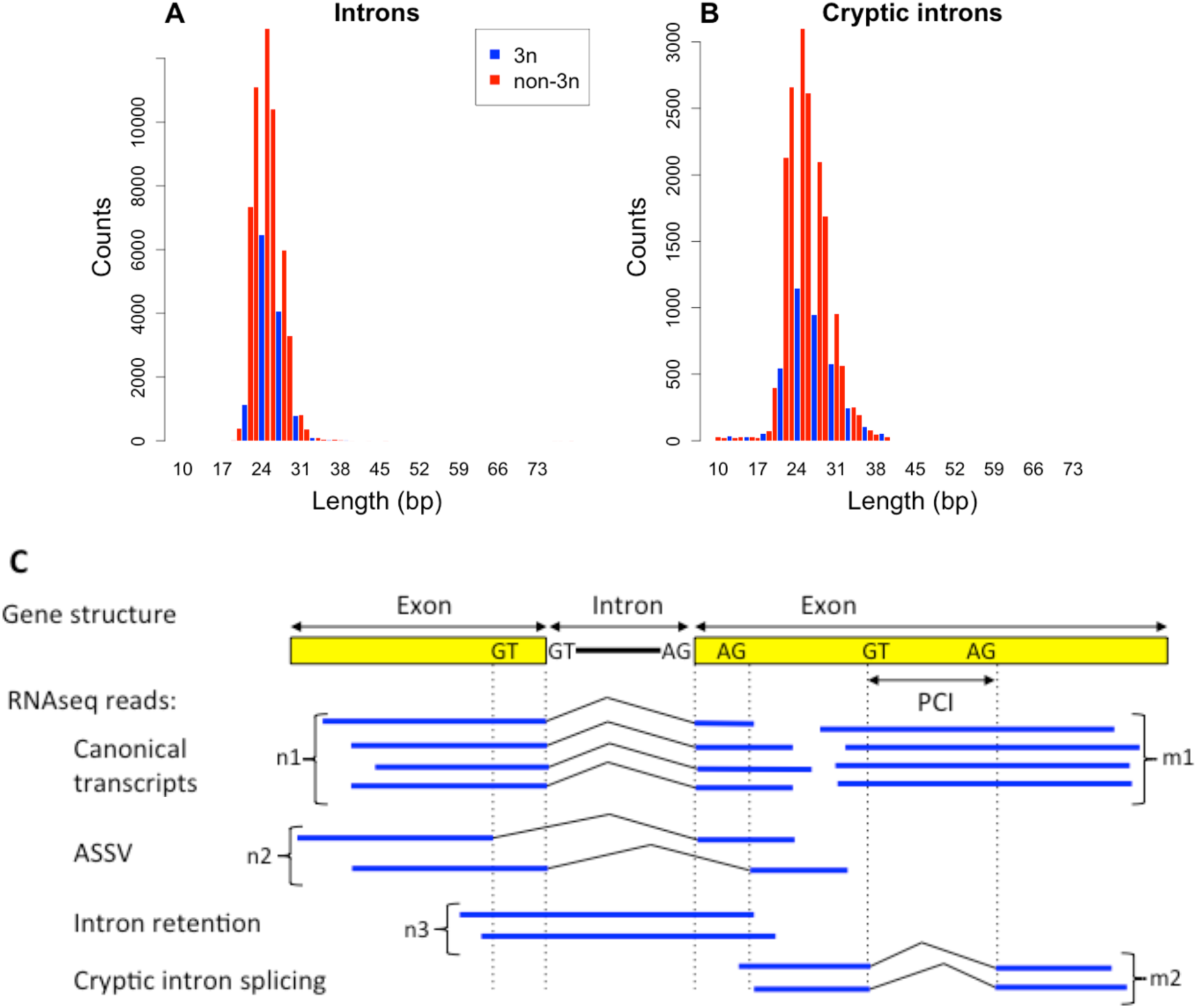
Introns and cryptic introns in Paramecium tetraurelia. (A) Length distribution of introns (N=65,159). (B) Length distribution of cryptic introns (N=20,719 cryptic introns detected in WT or NMD-deficient cells). Introns and cryptic introns of length multiple of three (3n), or non-multiple of three (non-3n) are displayed respectively in blue and red. (C) Quantification of splicing variation. For each intron, we identified all RNAseq reads spanning both flanking exons, and counted the number of reads corresponding to the canonical transcript (n1), to usage of 5' or 3’ alternative splice sites (ASSV, n2) and to intron retention (n3). The IR rate is defined as n3/(n1+n2+n3), the ASSV rate is n2/(n1+n2+n3). Similarly, for potential cryptic introns (PCIs), the splice rate is defined as m2/(m1+m2).

## RESULTS AND DISCUSSION

### Quantification of splicing variants in Paramecium

For a given gene, the abundance of splicing variants depends both on the intrinsic strength of splicing signals, and on the relative stability of the different variants. Thus, to study the determinants of alternative splicing in *Paramecium tetraurelia*, we sequenced the polyadenylated RNA fraction of cells, either in normal state (hereafter denoted wild-type, WT), or rendered NMD-deficient by knocking down one of the main components of the NMD machinery (Upf1, Upf2 or Upf3). The inactivation of Upf genes leads to stabilization of PTC-containing transcripts that would normally be degraded by the NMD machinery, thus providing a proxy for the intrinsic splicing efficiency of introns.

We generated 10 RNA-seq datasets (Supplemental Table S1): 6 distinct NMD knockdown experiments, and 4 replicates of WT cell cultures (see methods). All biological replicates gave similar results (Supplemental Figures S1 and S2). We therefore pooled the sequencing datasets, to increase the per gene read depth (50% of genes have a read depth >41 and >85 in WT and in NMD-deficient samples respectively). We detected splicing events by mapping sequence reads to the genome. These splicing events were then compared to gene models of the reference genome annotation, which includes 39,642 protein-coding genes, among which 31,632 contain introns (Aury et al. 2006).

We detected three types of AS events (Fig. 1C): intron retention (IR), alternative splice site variants (ASSV), and splicing of cryptic introns (i.e. introns with both splice sites located within an annotated coding exon). It is important to note that the classification of splice variants relies on the definition of a canonical form (Fig. 1C): the distinction between a “ cryptic intron “ and a “retained intron” depends on which variant is considered as the reference. For the vast majority of introns (97.8%), we observed one single major splice form, at least five times more abundant than other forms (Supplemental Fig. S3). We therefore decided to define the canonical form as the one that is the most abundant in WT cells (see Supplemental Text S1). To be able to identify canonical forms, we restricted all subsequent analyses to genomic segments covered by at least 10 RNAseq reads in WT samples. This subset includes 65,159 annotated introns (which constitute our reference intron dataset).

To compare alternative splicing rates between different samples, it is necessary to normalize variant counts by the sequencing depth (Kim et al. 2007). For introns, we computed the rates of retention and ASSV, defined as the proportion of variant reads among all reads spanning these reference introns (Fig. 1C). For cryptic introns, we considered all DNA segments potentially subject to cryptic splicing, i.e. segments of length 20 to 35 nt (matching the size distribution of observed introns and cryptic introns, Fig. 1), entirely located within an exon, starting with GT and ending with AG. These segments will hereafter be referred to as “ potential cryptic introns” (PCIs). The rate of cryptic intron splicing is defined by the proportion of spliced reads among all reads spanning PCIs (Fig. 1C).

The average IR rate is about 5 times higher than the ASSV rate, and 100 times higher than the splice rate of PCIs (Table 1). However, given the very large number of PCIs (on average there are 34.9 PCIs per gene, vs. only 2.3 introns), cryptic introns constitute a substantial fraction (6.9%) of all splice variants. Overall, combining all samples (WT and NMD-deficient), 95.0% of intron-containing genes show evidence of splicing variability in at least one of their introns, and 32.3% of genes contain at least one detected cryptic intron (Supplemental Table S2). IR and ASSV rates are comparable to those observed in humans (Table 1). We did not observe any case of exon skipping in paramecia, but we detected 20,719 cryptic introns, 20 times more than reported in *Arabidopsis thaliana* and in humans (Marquez et al. 2015). This probably reflects the fact that the splicing machinery of paramecium only recognizes very short introns, which increases the risk of excising cryptic introns within exons, but precludes exon skipping.

**Table 1.** Impact of NMD on steady-state levels of splice variants.

|  | *P. tetraurelia* | Human |
| --- | --- | --- |
| Number of protein-coding genes | 39,642 | 19,919 |
| Mean (median) number of introns per gene | 2.3 (2.0) | 9.3 (7.0) |
| Average ASSV rate per intron | 0.6% | 1.9% |
| Average IR rate per intron | 3.3% | 3.4% |
| Average splice rate per PCI | 0.026% | NA |

We classified splice variants in three categories according to their impact on the translation reading frame. Among all PCIs, 63.8% are predicted to lead to NMD-visible (i.e. PTC-containing) transcripts in case of splicing, while 80.1% of introns are predicted to be NMD-visible in case of retention. As expected, the abundance of NMD-visible variants is strongly increased in NMD-deficient cells compared to WT cells (Fig. 2). For NMD-invisible variants, we observed a weak but significant increase in NMD-deficient cells compared to WT cells (Fig. 2). This increase probably reflects an indirect consequence of NMD inactivation: in many species, genes encoding splicing factors are regulated by AS-NMD (Lareau et al. 2007; Ni et al. 2007), and we observed the same pattern in paramecia (Supplemental Text S2, Supplemental Fig. S4). Hence, the inactivation of NMD is expected to alter the efficiency of the splicing machinery, and thereby to indirectly affect the overall splicing pattern. The variation in AS rate for NMD-invisible variants is however much weaker than that observed for NMD-visible variants, which indicates that NMD directly affects the steady state levels of PTC-containing splice variants.

**Figure 2:**
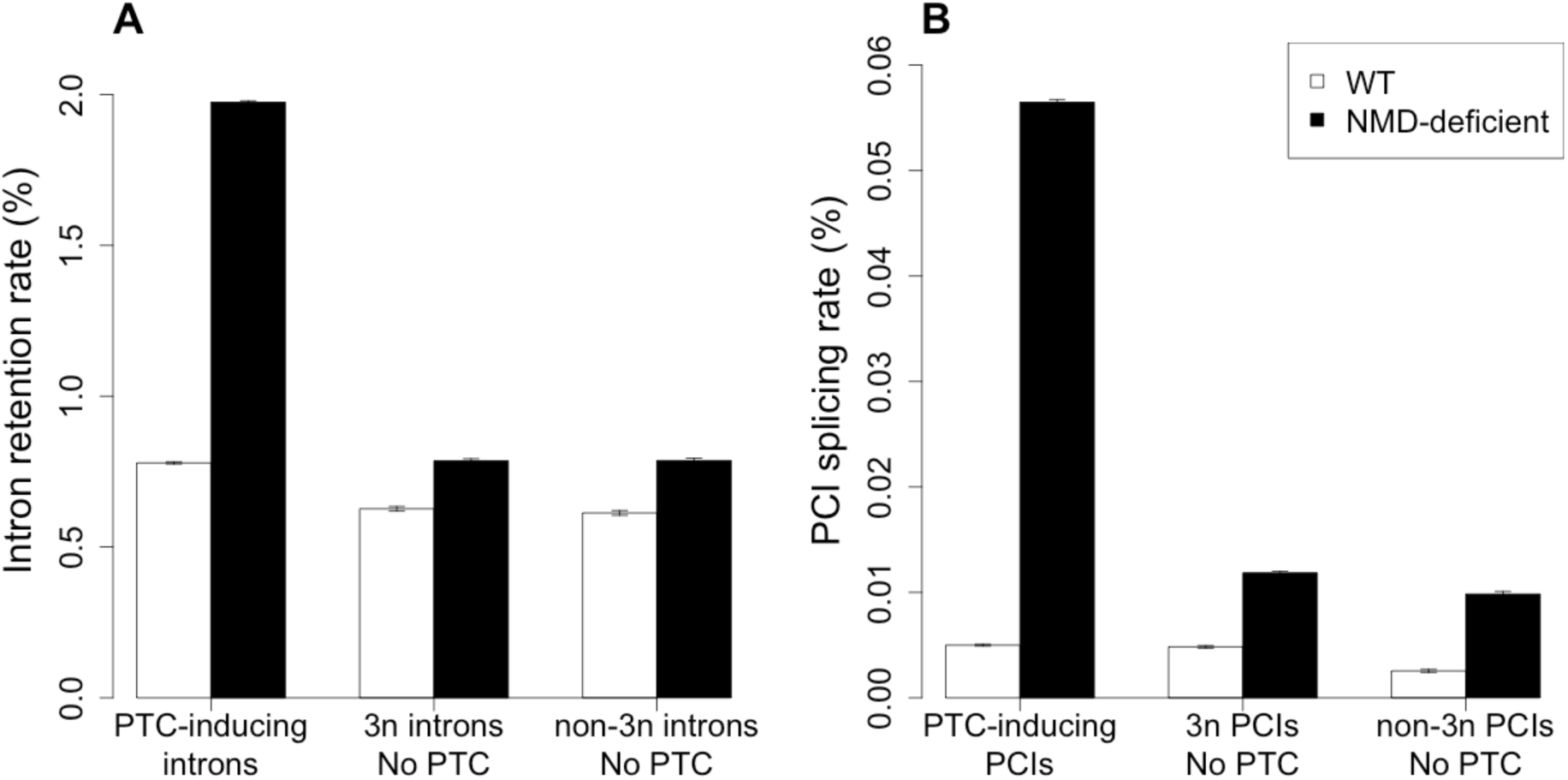
Impact of NMD on observed AS rates. AS events (IR or cryptic intron splicing) are classified into three groups according to their NMD-visibility: PTC-inducing events (i.e. NMD-visible); events that do not introduce frameshift or PTC (3n no PTC); events that create a frameshift but without introducing a PTC (non-3n no PTC). The two latter categories are not detectable by NMD. AS rates in WT and in NMD-deficient cells were computed globally within each bin, as the proportion of AS reads among all reads spanning introns (or PCIs) from that bin. Error bars represent the 95% confidence interval of this proportion. (A) Intron retention (N= 65,159 introns). (B) Splicing of PCIs (N= 1,383,067 PCIs).

### Lower rate of alternative splicing in long and highly expressed genes

The previous observations indicate that AS-NMD might potentially contribute to the post-transcriptional regulation of many genes. However, they are also compatible with the hypothesis that most splice variants are errors, and that NMD is used as a surveillance mechanism to degrade erroneous transcripts. This “ noisy splicing” model makes several testable predictions, which are based on three points. First, the cost of splicing errors is expected to increase with gene expression level: for a given splicing error rate, the waste of resources (both in terms of metabolic cost and of futile mobilization of cellular machineries) will be larger for highly expressed genes, and hence, the selective pressure on splicing accuracy is expected to be stronger. In other words, if AS events predominantly correspond to errors, the selection-mutation-drift theory predicts that the AS rate should correlate negatively with gene expression level. To test this prediction, we classified introns (or PCIs) into 10 bins of equal sample size according to their gene expression level, and computed the AS rate within each bin. In agreement with the noisy splicing model, we observed a strong decrease in AS rate with increasing expression level, for IR (Fig. 3A), ASSV (Fig. 3B) and cryptic intron splicing (Fig. 3C). This pattern is observed in both WT and NMD-deficient cells, which indicates that the observed variations reflect differences in intrinsic splicing efficiency.

The second point is that, for a given splicing error rate per intron, the rate of production of spurious transcripts increases with the number of introns present in a gene: the greater the number of introns, the greater the risk of having at least one error. The selective pressure on the strength of splice signals of each intron is therefore expected to increase with the number of introns in a gene, and hence the AS rate (per intron) should be lower in genes with more introns. To test this prediction, we classified introns into 3 groups according to the number of introns present in their gene: genes with 1 intron, with 2 to 3 introns, and with at least 4 introns (mean = 5.2 introns) (the three groups correspond respectively to 27.8%, 43.0% and 29.2% of intron-containing genes). We then binned each group according to gene expression level and computed the AS rate per bin. Again, observations perfectly match predictions: for a given expression level, the AS rate per intron is higher in genes with fewer introns, both for IR (Fig. 3D) and for ASSV (Fig. 3E).

**Figure 3:**
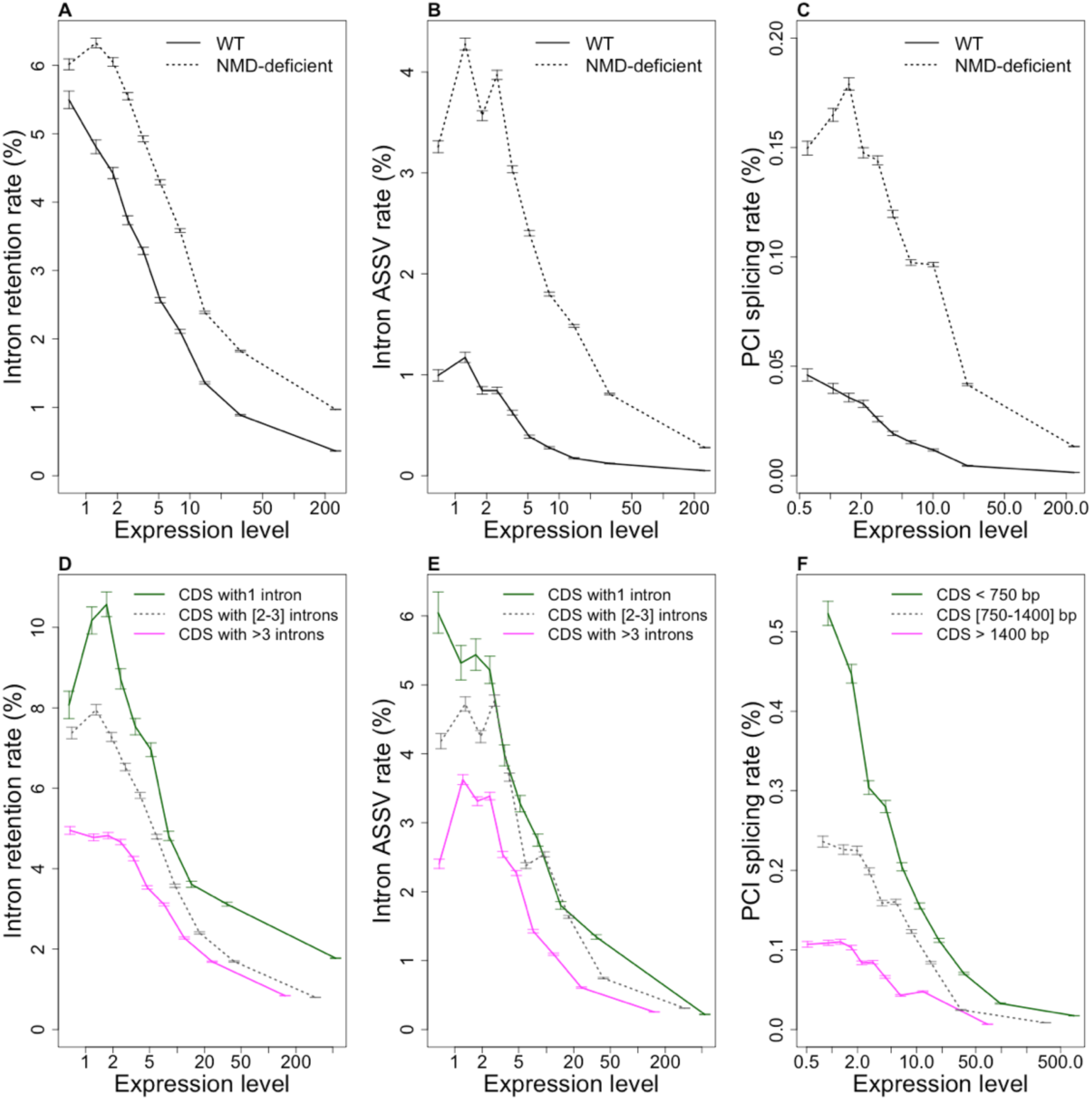
Relationship between AS rate and gene features: expression level, number of introns or length of coding regions. Introns (N= 65,159) and PCIs (N= 1,383,067) were classified into ten bins of equal sample size, according to gene expression levels in WT cells. The AS rate was computed globally within each bin, as the proportion of AS reads among all reads spanning introns (or PCIs) from that bin. Error bars represent the 95% confidence interval of this proportion. (4) IR rate; (B) 455V rate; (C) Rate of splicing at potential cryptic introns. (D,E) same as (4,B), but introns were first classified into three bins, according to the number of introns of the gene in which they are located: genes with 1 intron (N=5,606 introns), genes with 2 to 3 introns (N=24,452 introns), genes with > 3 introns (N=35,101 introns). (F) same as (C), but PCIs were first classified into three bins, according to the length of the coding region (CD5) in which they are located: CD5 < 750 bp (N=169,030 PCIs), CD5 from 750 to 1400 bp (N=406,460 PCIs), CD5 > 1400 bp (N=807,577 PCIs). (4,B,C): 45 rates were measured in normal cells (WT, black line) and in NMD-deficient cells (dashed line). (D,E,F): 45 rates were measured in NMD-deficient cells. Expression levels (RPKM) are represented in log scale.

The third point is that the risk of cryptic intron splicing increases with the number of PCIs, and therefore with the length of coding sequences (CDSs). The selective pressure to limit the strength of cryptic splice signals should therefore increase with CDS length, and PCIs in long CDSs should have a lower splicing rate compared to PCIs in short CDSs. To test this prediction, we classified PCIs into 3 groups according to the length of the CDS in which they are located (each group corresponds to 1/3 of all genes), and then binned each group by gene expression level and computed the PCI splicing rate per bin. Again, the predictions of the model fit the observations: for a given expression level, the splicing rate per PCI is lower in genes with longer CDSs (Fig. 3F). Thus, all observations fit the three predictions of the “ noisy splicing” model.

### The genome-wide AS pattern is dominated by splicing errors

The previous results indicate that the level of constraints against splicing errors is maximal in highly-expressed genes containing many introns and/or encoding long CDSs (Fig. 3). The strong relationship between AS rate and expression level can be used to quantify the splicing error rate in each bin of expression. The proportion of AS events that correspond to splice errors *(P^e^)* is given by:

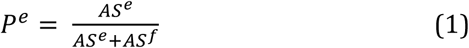

where *AS^f^* is the rate of functional AS events and *AS^e^* is the rate of erroneous splicing.

The ratio of the AS rate in a given bin of expression (*i*) over the AS rate in highly constrained genes (*h*) is given by:

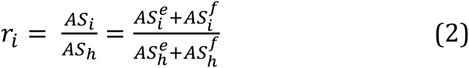

Under the assumption that the rate of functional AS events is the same for both gene classes (*AS_h_^f^ = AS_i_^f^* = *AS^f^*), the proportion of splicing errors in expression bin (*i*) can be written as:

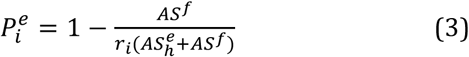

If selection is very strong in the set of highly constrained genes, so that the splicing error rate is negligible compared to the rate of functional AS events in that gene set (i.e. *AS_h_^e^* ≪ *AS^f^*), then equation (3) simplifies to:

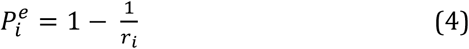

As a reference for highly constrained genes we considered genes with a high expression level (top 10%) and with more than 3 introns (for the quantification of erroneous ASSV and IR events) or with a CDS longer than 1400 bp (for the quantification of erroneous cryptic intron splicing). In WT cells, we observed that the ratio of the AS rate in genes with median expression level over the AS rate in highly constrained genes are respectively *r_i_*=12.0, *r_i_*=20.3 and *r_i_=*49.3 for IR, ASSV and cryptic intron splicing. According to equation (4), this implies that for a median gene, 92% to 98% of splice variants detected in WT cells result from errors, and this proportion might even be higher if the splicing error rate in highly expressed genes is not negligible (equation 3).

These estimates are based on the assumption that on average, the rate of functional AS does not vary with gene expression level (i.e. *AS^h^_f_= AS^l^_f_ = AS_f_* in equation 3). One may argue however that variation in AS rate with expression level might reflect differences in the propensity to use AS-NMD: it is in principle possible that weakly expressed genes are more prone to use AS-NMD to fine-tune their expression level (i.e. *AS^l^_f_* > *AS^h^_f_*). For instance, one might speculate that highly expressed genes are preferentially regulated at the transcriptional level, to avoid the waste of resources caused by the posttranscriptional AS-NMD pathway.

To test this hypothesis, we analyzed splicing variants according to their NMD-visibility. We observed a strong negative relationship between AS rate and gene expression level, both for NMD-visible and NMD-invisible splicing variants (Fig. 4 for WT cells and Supplemental Fig. S5 for NMD-deficient cells). In other words, weakly expressed genes show a high rate of alternative splicing events, even for NMD-invisible splicing events, which, by definition, cannot contribute to the regulation of gene expression by AS-NMD. Thus, the observed relationship between AS rate and gene expression level cannot be explained by a higher propensity of weakly expressed genes to be regulated by AS-NMD. The most parsimonious explanation is that the excess of AS in weakly expressed genes compared to highly expressed genes simply reflects differences in the selection-mutation-drift equilibrium: these genes are under weaker selective pressure for splicing accuracy, and hence show a higher rate of splicing error. If this interpretation is correct, then our calculations imply that for a median gene, at least 92% to 98% of splice variants detected in WT cells correspond to weakly deleterious errors.

**Figure 4:**
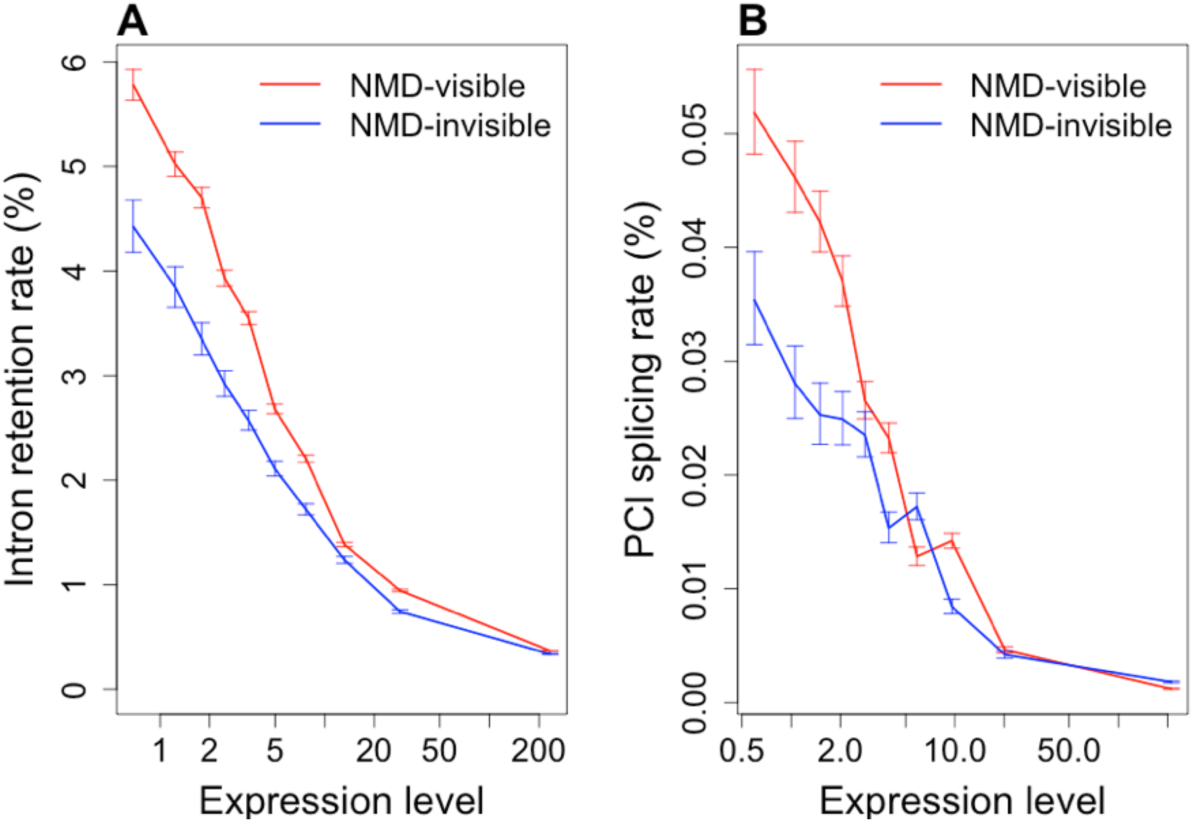
Relationship between AS rate and expression level, for NMD-visible or NMD-invisible AS events. (A) Introns were first classified into two groups according to their NMD-visibility in case of retention events (N= 52,163 NMD-visible introns, in red, and N=12,996 NMD-invisible introns, in blue), and then further grouped into ten bins of equal sample size, according to gene expression levels in WT cells. IR rates (in WT cells) were measured globally in each bin. Error bars represent the 95% confidence interval of the proportion of AS reads. (B) Same as (A), but for the splicing of PCIs: N= 882,579 NMD-visible PCIs, and N=500,488 NMD-invisible PCIs. Expression levels (RPKM) are represented in log scale.

### A dual strategy to limit the cost of splicing errors

In NMD-deficient cells, the IR rate is much higher for NMD-visible introns than for NMD-invisible introns, which indicates that the former have a lower intrinsic splicing efficiency (Fig. 2A). The difference in intrinsic splicing efficiency results, at least in part, from a difference in the strength of splice signals: on average, 77.4% of NMD-invisible introns match the consensus splicing signals [GTA..TAG], compared to only 69.8% for NMD-visible introns (Chi-squared test = 289.1, p < 10^-15^). However, in WT cells, the observed AS rate is similar for both categories of introns. This implies that the efficacy of NMD to eliminate transcripts with retained introns is strong enough to compensate the lower intrinsic splicing efficiency of NMD-visible introns.

The same pattern is observed for PCIs: in WT cells, NMD-visible and NMD-invisible PCIs show similar rates of splicing (Fig. 2B, Supplemental Fig. S6A), despite the fact that the intrinsic rate of splicing of PCIs (observed in NMD-deficient cells) is about 5 times higher for NMD-invisible compared to NMD-visible PCIs (Fig. 2B, Supplemental Fig. S6B). Thus, again, the higher intrinsic propensity of NMD-visible PCIs to be spliced out is compensated by the activity of NMD in WT cells.

### What is true for paramecia is true for humans

To test whether the observations that we made in a unicellular organism (*P. tetraurelia*) hold true in multicellular eukaryotes, we quantified ASSV in human introns, using previously published RNAseq datasets coming from 25 different tissues or cell types (Supplemental Table S3). We also re-analyzed a dataset published by Braunschweig et al. (2014), which provides a quantification of IR rates of human introns in 52 different tissues and cell types. In agreement with previous reports (Braunschweig et al. 2014), we observed that the IR rate (averaged over the 52 samples) decreases with increasing gene expression level. According to the authors, this observation supports their conclusion that gene expression is regulated through NMD acting on transcripts with retained introns (Braunschweig et al. 2014). However, the negative relationship between IR rate and expression level is observed both for NMD-visible events and for NMD-invisible events (Supplemental Fig. S7A), which is not consistent with the AS-NMD model. Moreover, we observed that for a given expression level, the IR rate (per intron) decreases with increasing number of introns in the gene (Fig. 5A). We observed exactly the same patterns for ASSV rates (Fig. 5B and Supplemental Fig. S7B). Thus, in humans as in paramecia, variations in ASSV and IR rates fit with the prediction of the noisy splicing model.

**Figure 5:**
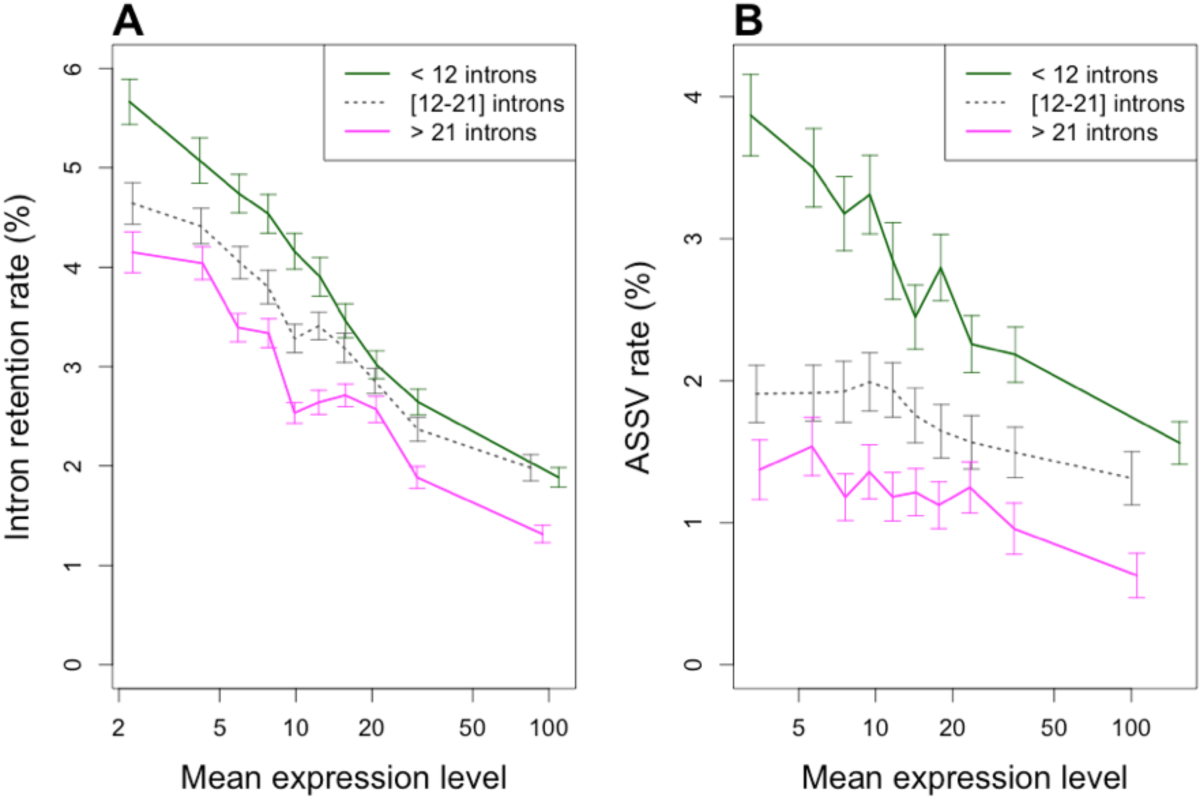
Relationship between AS rate, expression level and number of introns in human genes. (A) IR rate (N=118,703 introns). (B) ASSV rate (N=102,697 introns). In both panels, introns were first classified into three groups of equal sample size, according to the number of introns of the genes in which they are located (genes with < 12 introns, genes with 12 to 21 introns, genes with > 21 introns), and then further grouped into ten bins of equal sample size, according to gene expression levels. We computed the average AS rate (IR or ASSV) over all introns within each bin. Error bars represent the 95% confidence interval of the mean. Expression levels (RPKM, averaged over the 52 samples) are represented in log scale

As a reference dataset of highly constrained human genes, we considered genes with a high expression level (top 10%) and with more than 21 introns (top 33%). The ratio of the AS rate in genes with median expression level over the AS rate in highly constrained genes are respectively *r_i_=*3.1 and *r_i_*=3.6 for IR and ASSV. According to equation (4), this implies that for median genes, at least 68% of ASSV events and 72% of IR events correspond to errors. It should be stressed that these estimates are based on the assumption that the error rate in the set of highly constrained genes is negligible. In paramecia, AS rates tend to plateau at high expression levels (Fig. 3), which is compatible with the hypothesis that this basal rate might correspond to functional splice variants. However, in human, contrary to paramecia, there is no sign that AS rates reach a basal value at high expression levels (Fig. 5). It is therefore likely that the splicing error rate is substantial, even in the reference dataset of highly constrained genes. Hence, the above estimates are certainly an underestimate of the true splicing error rate in humans.

### Fitness impact of mis-splicing in humans

One strong assumption of the noisy splicing model is that the fitness impact of splicing errors increases with expression level. To test this hypothesis, we analyzed patterns of polymorphism in the vicinity of human splice sites. Splicing imposes strong constraints on donor and acceptor sites (defined as the first and last 2 nt of introns): 99.1% of human introns start with GT, and 99.8% end with AG. As expected, these sites show evidence of strong purifying selection: the SNP density is 4.5 fold lower at splice sites than in flanking third codon positions (Fig. 6A). We quantified this selective pressure by measuring the ratio π_spl_/π_3_, where π_spl_ is the SNP density at splice sites, and π_3_ the SNP density at flanking third codon positions. We binned introns by gene expression level, and computed this ratio in each bin. Interestingly, the π_spl_ /π_3_ ratio is strongly correlated to gene expression level (R^2^=0.89, p<10^-9^), with a five-fold difference between lowly and highly expressed gene sets (Fig. 6B, black dots). It should be stressed that the fraction of introns matching the GT..AG consensus does not vary with gene expression level (Fig. 6D). This implies that mutations occurring at donor and acceptor sites are ultimately counter-selected, even in weakly expressed genes. However, our observations (Fig. 6B) show that these mutations are more rapidly purged in highly expressed genes. This demonstrates that the fitness cost of mis-splicing increases with gene expression level.

**Figure 6:**
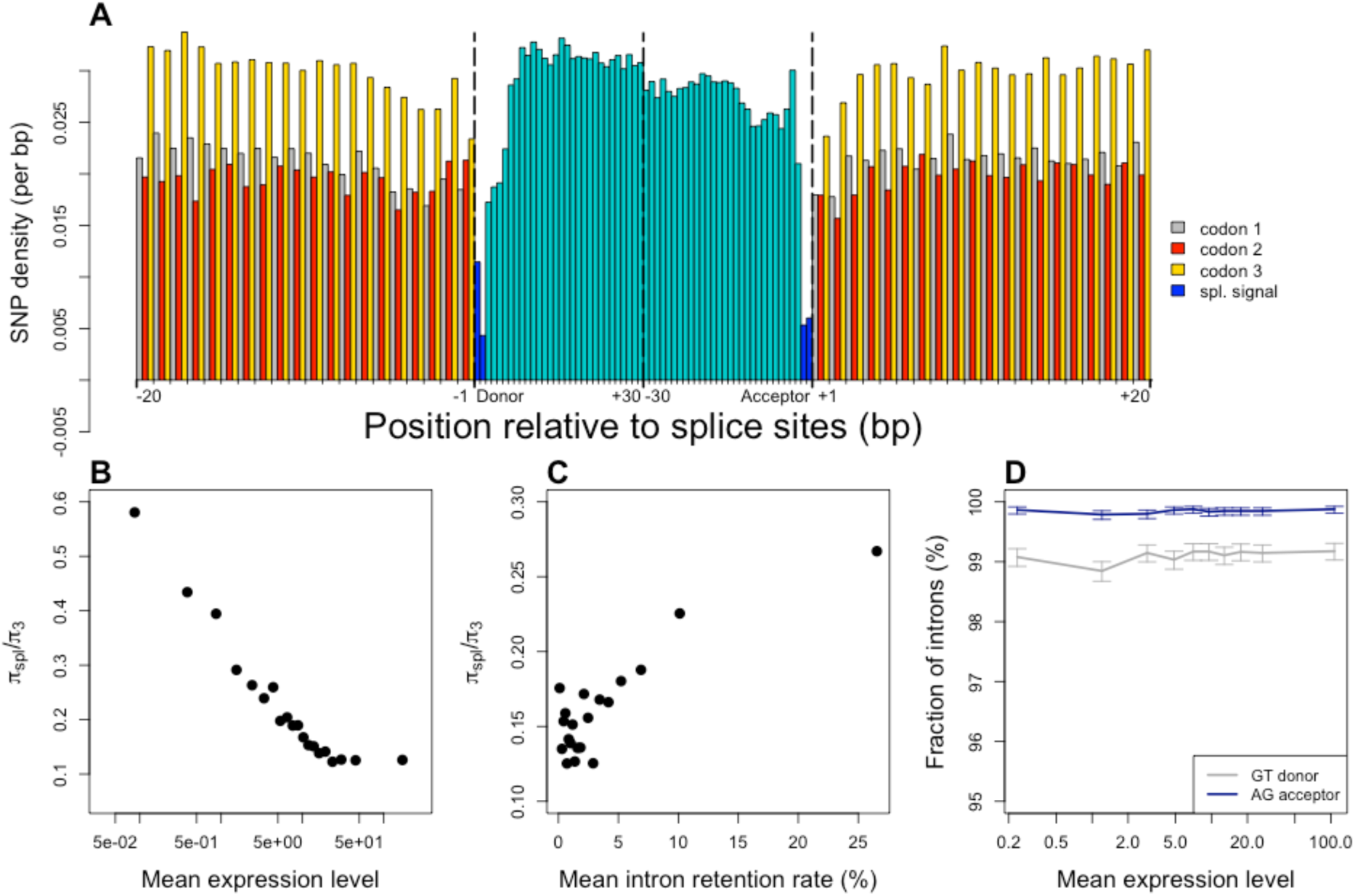
Variation in selective constraints on splice signals in human genes. (A) SNP density was measured in the vicinity of exon-intron boundaries (first and last 30 bp of introns, and 20 bp of flanking exons), over all introns located between coding exons (N=170,015). Splice sites (first and last 2 bp of introns) are displayed in dark blue, other intron positions in light blue. Within coding regions, the SNP density at each site was computed separately for the three codon positions (grey: position 1, red: position 2, yellow: position 3). (B) The level of selective constraints on splice signals increases with gene expression level. Introns were classified into bins of equal sample size, according to gene expression levels. Within each bin, the fitness impact of mutations on splice sites was estimated by measuring the ratio *π_spι_/π*_3_, where π_spι_ is the SNP density at splice sites, and π_3_ the SNP density at flanking third codon positions. (C) The level of selective constraints on splice signals decreases with increasing IR rate. Introns were classified into bins of equal sample size according to their average retention rate, and the ratio *π_spι_/π_3_* was measured in each bin. (D) The fraction of introns with consensus splice signals does not vary with gene expression level. The proportion of introns matching the consensus splice donor (GT), and the proportion of introns matching the consensus splice acceptor (AG) was computed for each bin of expression level. Error bars represent the 95% confidence interval of this proportion. (B and D): mean expression levels (RPKM) are represented in log scale.

To test whether IR rate co-varies with the fitness impact of mis-splicing, we binned introns according to their IR rate, and computed π_spl_/π_3_ in each bin. We observed a positive correlation between π_spl_/π_3_ and the average IR rate per bin (R^2^=0.76, p<10^-6^), with a twofold increase between bins of low IR compared to bins of high IR (Fig. 6C). Again, it is important to stress that the frequency of introns matching the GT..AG consensus does not vary with IR rate (Supplemental Fig. S8). This implies that mis-splicing is deleterious, even in introns with high IR rate. However, in agreement with the noisy splicing model, introns that show a high IR rate correspond to introns where mis-splicing is relatively less deleterious.

## CONCLUSION

The efficiency of excision of introns by the spliceosome is affected by different signals, located within introns and flanking exons (splice sites, branch point, polypyrimidine tract, splicing enhancers or silencers). Besides the two splice sites that are critical for the splicing reaction (almost always GT for the donor and AG for the acceptor), all other signals tolerate some sequence flexibility. The probability for a mutation affecting a splicing signal to reach fixation depends on its fitness impact (i.e. the selection coefficient, *s*) and on the power of random genetic drift (i.e. the effective population size, *N_e_*). There is therefore necessarily a limit to the point up to which selection can optimize the strength of splice signals: if the splicing error rate is already low, any mutation that further improves splicing efficiency will necessarily have a weak fitness impact, and hence will be subject to random drift (the so-called drift barrier effect (Sung et al. 2012a)). This drift barrier therefore determines a basal splicing error rate, which depends on the mutation rate, on *N_e_* and on the fitness cost of splicing errors (*s*).

For a given error rate, errors are expected to be more costly (in terms of metabolic resources and mobilization of cellular machineries) in highly expressed genes. Hence the fitness cost of mis-splicing is expected to increase with increasing expression level. Indeed, this is precisely what we observed in humans: the strength of selection against deleterious mutations at splice sites is strongly correlated to gene expression level (Fig. 6B). Since the risk of producing erroneous transcripts increases with the number of introns, this implies that all else being equal, there should be a stronger selective pressure against mis-splicing in intron-rich genes. The mutation-selection-drift theory therefore predicts that introns from weakly expressed/intron poor genes should accumulate more non-optimal substitutions in their splice signals, and therefore should show a higher splicing error rate. The relationships that we observe between AS rate, expression level and intron number are perfectly consistent with these predictions, both in human (Fig 6) and in paramecia (Fig 3).

There are two possible ways to limit the deleterious impact of erroneous splicing: either i) improve the strength of splicing signals to increase intrinsic splicing efficiency and avoid the use of cryptic signals (error prevention), or ii) ensure that transcripts are degraded by NMD in case of splicing error (error mitigation). We observed that both strategies are used: there is a deficit of introns and cryptic introns that cannot trigger NMD in case of splicing error, and the rare introns that are not NMD-visible show stronger splicing signals (Supplemental Text S3, Supplemental Fig. S9). The analysis of AS rate in NMD-deficient cells shows that NMD-invisible introns have a much higher intrinsic splicing accuracy than NMD-visible ones. This difference demonstrates that the biophysical limits of splicing accuracy have not been reached, and that it would be possible to further improve splicing accuracy of NMD-visible introns by genetic engineering. However, the mutation-selection-drift theory predicts that once the basal splicing error rate has been reached, by error prevention or by error mitigation, then selection cannot further improve splicing efficiency. Thus, this model predicts that the steady state level of erroneous transcripts (after quality control by NMD) should be the same for NMD-visible and NMD-invisible introns. And this is precisely what we observed: in WT cells, NMD-visible and NMD-invisible AS events show similar rates (Fig. 2).

The fitness cost of splicing errors depends on the frequency of transcripts subject to at least one erroneous splicing event. Owing to the short length of RNAseq sequence reads, it is not possible to directly quantify AS rates per transcript. However, given that AS rates (per intron) are similar in human and in paramecia (Table 1), and that human genes contain on average 3 to 4 times more introns than paramecia, this implies that the frequency of transcripts subject to at least one erroneous splicing event must be much higher in human than in paramecia. This is consistent with the drift-barrier hypothesis, which predicts that humans should have a higher splicing error rate (per gene), owing to their larger mutational targets (more introns) and to their smaller effective population size (Sung et al. 2012a, 2012b).

There is clear evidence that some AS events are functional (Kelemen et al. 2013). Notably we observed that AS-NMD probably plays an important role in the regulation of genes encoding splicing factors in paramecia (Supplemental Text S3), as previously shown in other eukaryotes (Lareau et al. 2007; Ni et al. 2007). However, AS-NMD cannot explain the strong relationship between AS rate and expression level that is observed for NMD-invisible splicing variants (Fig. 4, Fig. 5A). It has been recently shown that the retention of introns in nuclear transcripts (the so-called detained introns) might also contribute to the regulation of gene expression, independently of NMD (Boutz et al. 2014). If weakly expressed genes were more prone to use this regulatory pathway, this might explain the relationship observed between expression level and IR rate. However, this model does not explain the relationship between IR rate and intron number (Fig. 3D, Fig. 5B), and most importantly, cannot explain the relationship between expression level and other classes of AS events (ASSV or cryptic intron splicing; Fig. 3, Fig. 5). The most parsimonious explanation is that the excess of AS in weakly expressed/intron-poor genes results from the accumulation of maladaptive substitutions, driven by random genetic drift in genes where the selective pressure is weaker. Our observations indicate that for median genes, the vast majority of observed splice variants correspond to errors, in contradiction with the panglossian view of a widespread role of AS-NMD in fine-tuning the expression of genes. Of course, this does not negate the importance of AS-NMD in the regulation of some genes. However, our results highlight the necessity of a careful consideration of non-adaptive hypotheses before concluding about the functionality of AS events.

## METHODS

### *Paramecium* strain, cell culture and inactivation of NMD

The entirely homozygous strain 51 of *P. tetraurelia* was grown in a wheat grass powder infusion medium bacterized with *Klebsiella pneumoniae* the day before use, and supplemented with 0.8 mg.l^-1^ β-sitosterol. NMD was inactivated either by RNAi-mediated silencing of UPF genes during vegetative growth of wild-type cells, or by generating somatic knockouts, *i. e.* clones in which these genes are deleted from the macronucleus. RNAi treatment was based on the doublestranded RNA feeding technique (Beisson et al. 2010): briefly, cells were fed for 7 days with *E. coli* (HT115) producing double-stranded RNA homologous to the target gene. Sequences used for silencing of UPF1A, UPF1B, UPF2, UPF3 and ICL7a (which encodes a cytoskeletal protein), were segments 1,885-2,289, 1,887-2,285, 1,143- 1,546, 18-422 and 1-580 of the genes (from the ATG), respectively. These genes can be accessed with ParameciumDB (http://paramecium.cgm.cnrs-gif.fr/) under accession numbers GSPATG00034062001, GSPATG00037251001, GSPATG00017015001, GSPATG00001393001 and GSPATG00021610001, respectively. Somatic knockouts were generated by applying RNAi treatment during the development of a new somatic macronucleus, which results in the deletion of the targeted genes (Garnier et al. 2004; Dubois et al. 2017): wild-type conjugating pairs were transferred to “ UPF” RNAi medium and, following their separation, individual exconjugants were isolated in the same medium. After 24 hours of growth, cells were transferred to standard growth medium. Among the viable exconjugants obtained, somatic UPF deletions were screened for based on the slow growth phenotype and the inability to undergo autogamy, and later confirmed by Southern blots and PCR (Supplemental Fig. S10).

### RNA-seq

Total RNA was extracted from cells grown on *K. pneumoniae* or the relevant feeding *E. coli* strains with the TRIzol (Invitrogen) procedure, modified by the addition of glass beads. For the first 4 RNA-seq datasets in Supplemental Table S1, poly(A) RNAs were purified from 100 μg of total RNA with the MicroPoly(A)purist kit (Ambion). 25% of the output was used for mRNA reverse transcription, using the SuperScript III kit (Invitrogen) and the anchor-oligo(dT) primer 5’-GCCCACCAGAGCCGGCGGATTTTTTTTTTTTTTTTT-3’. After alkaline lysis of RNA and removal of the oligo(dT) primer with G-50 columns (GE Healthcare), a poly(G) tail was added to singlestranded cDNAs with terminal transferase (NEB) following the producer’s instructions. After phenol purification and ethanol precipitation, cDNAs were made double-stranded using the Phusion PCR enzyme (Finnzymes) and the anchor-oligo(dC) primer 5’-GCCCACCAGAGCCGGCGGACCCCCCCCCCCCCCCCC-3’. Double-stranded DNA was then purified using the Qiagen PCR purification kit, and cDNA libraries were amplified by 15 cycles of PCR with the anchor primer. cDNA libraries were digested by EciI restriction enzyme (NEB) and purified (Qiagen) before addition of Illumina adaptors. For the last 6 RNA-Seq datasets, library preparation and Illumina sequencing were performed at the ENS Genomic Platform (Paris, France). Poly(A) RNAs were purified from 1 μg of total RNA using oligo(dT). Libraries were prepared using the strand non-specific RNA-Seq library preparation TruSeq RNA Sample Prep kit (Illumina), and multiplexed by 3 on 2 flowcell lanes. 101-bp paired-end read sequencing was performed on a HiSeq 1500 device (Illumina).

### Read mapping

The sequencing of these 10 samples yielded a total of 40.8 Gb (from 247,653,027 fragments), 25.1 Gb from NMD-deficient cells, and 15.7 Gb from control cells (Table S1). Reads were mapped against the *P. tetraurelia* reference genome assembly (accession number: CAAL01000000) (Aury et al. 2006), using TopHat (version 1.4.1) (Trapnell et al. 2009). The minimal and maximal intron lengths were set to 10 nt and 500000 nt respectively. Reads that mapped at multiple positions on the genome were excluded from further analyses. Read coverage along transcription unit was obtained using annotated gene models from the reference genome (Aury et al. 2006). The expression level of genes was measured in reads per kilobase per million mapped reads (RPKM).

### Detection of splicing events

For each annotated intron, we counted the number of mapped reads spanning both extremities (Fig. 1C). Reads aligning to the genome sequence without any gap were counted as intron retention. Reads showing a deletion corresponding exactly to the annotated intron were counted as splice events. Reads with a deletion that does not match the annotated intron (at one or both extremities) were counted as alternative splice site variant (ASSV). Reads showing a deletion entirely located within an annotated coding exon were counted as cryptic intron splicing events (Fig. 1C).

The cell cultures that we analyzed are totally homozygous. However it is important to note that in paramecia, the macronuclear genome is highly polyploid, and that the different copies of a same gene may differ due to heterogeneity in the process of excision of internal eliminated sequences (IESs) (Duret et al. 2008). Thus, a fraction of the diversity detected in the transcriptome may in fact result from this macronuclear genomic heterogeneity. Among all alignment gaps detected by TopHat, more than 97% match the consensus intron boundaries (GT/AG), which indicates that most of them correspond to *bona fide* splice events. To avoid any confusion between splice variants and IES excision variants, we counted as splice variants only those matching the GT/AG consensus.

The classification of splice variants (IR, ASSV or cryptic intron splicing) was based on the comparison with the canonical form, defined as the major form observed in WT cells. Among the 90,287 annotated introns, we selected those that are spanned by at least 10 reads in WT samples (N=70,242). Among those ones, 4,045 were never observed as spliced and 1,038 correspond to minor splice forms. Thus, our reference data set includes 65,159 introns (72% of the initial data set).

### Quantification of AS rate

One important goal of this study was to analyze the relationship between AS rate and gene expression level. The AS rate at a given intron is defined by the proportion of splice variant reads among all reads spanning that intron (Fig. 1C). One difficulty is that the precision of this metric is strongly dependent on the sequencing read depth, and hence, the measure of the AS rate is much less accurate in weakly than in highly expressed genes. To circumvent this problem, we binned introns (or PCIs) by expression level, and then measured the global AS rates in each bin (defined by the proportion of splice variant reads among all reads in that bin).

### Analysis of intron retention in humans

Braunschweig et al (2014) analyzed 52 RNAseq samples from different tissues and cell types to quantify intron retention in human genes. For each gene, they selected one representative transcript, based on Ensembl annotations. Their initial dataset includes 202,973 introns from 20,959 protein-coding genes (Supplemental Tables S6 and S8 from Braunschweig et al (2014)). We computed the average gene expression level of each gene over the 52 samples, using data provided by the authors. We excluded data from genes that are not mapped on chromosomes of the reference genome assembly (N=18,546 introns from 2,185 genes annotated on unmapped contigs or additional haplotypes), or for which expression data were not available (N=4,844 introns from 871 genes).

To analyze the AS rate according to NMD visibility we also excluded from their dataset all introns located within UTRs or within truncated CDS (i.e. CDS lacking start or stop codon, or containing an internal stop codon): N= 10,780 introns from 912 genes. The final dataset includes introns from 16,991 genes.

For each intron, Braunschweig et al (2014)quantified retention rates in all samples where it showed sufficient read depth (> 10 reads spanning each flanking exon boundary). Among the 170,015 introns, we excluded those corresponding to minor splice forms (i.e. with an IR rate ≥ 50%, N=580 introns), and selected all those for which the retention rate had been quantified in at least 10 samples. For each of the selected introns (N=118,703), we computed the average retention rate over all available samples (median = 38 samples).

### Analysis of ASSV in humans

We estimated ASSV frequencies in 25 human tissues and cell lines, using 110 publically available RNA-seq samples (Supplemental Table S3), corresponding to a representative subset of the samples analyzed by Braunschweig et al (2014). To increase comparability among samples, for paired-end data we analyzed only the first read of the pair and stranded samples were treated as unstranded. We aligned the RNA-seq data on the human genome (hg38 assembly, downloaded from Ensembl release 84) using TopHat 2.0.4 with the following options: minimum intron size for junction discovery = 40 nucleotides (nt), maximum intron size = 1 million nt, maximum 1 mismatch per read segment, anchor size 8 nt, no mismatches allowed in the anchor region, no coverage search. To aid the spliced read mapping process, we provided as an input for TopHat the set of introns annotated in Ensembl release 84, with the −j option. We re-estimated the splice junction frequencies using uniquely mapping reads, annotated with the NH:i:1 tag in the original TopHat alignments. For each tissue/cell line, we combined read counts from all available samples.

For each intron from Braunschweig dataset (see above), we evaluated whether its 5’ or 3’ splice site were connected with alternative splice sites. We note E1 and E2 the annotated splice sites that border the intron, in 5’-3’ orientation. In a given tissue (*i*), we note *nE1E2i* the number of spliced reads corresponding to the annotated splicing event, *nE1E2_i_* the number of spliced reads that connect other 5’ splice sites of the same gene with the 3’ splice site E2, and *nElEa_i_* the number of spliced reads that connect the 5’ splice site E1 with other 3’ splice sites of the same gene. We then computed the alternative splice site variant frequency:

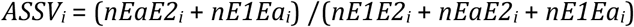

For a given intron, this parameter was computed only in tissues with sufficient read depth ((*nE1E2_i_* + *nEaE2_i_* + *nElEa_i_*) > 10 reads). We excluded 3,075 introns corresponding to minor splice forms (i.e. mean ASSV rate ≥ 50%), and selected all introns for which the ASSV rate had been quantified in at least 10 tissues. For each of the selected introns (N= 102,697), we computed the average ASSV rate over all available samples (median = 22 samples).

### Definition of NMD-invisible alternative splicing events in humans

In mammals, NMD is able to recognize and degrade PTC-containing transcripts only if the PTC occurs more than 50 nucleotides upstream of the last exon-exon junction (Popp and Maquat 2013; Lindeboom et al. 2016). Hence alternative splicing events (IR or ASSV) affecting last introns were classified as NMD-invisible, whereas the other were classified as potentially NMD-visible.

### Analysis of polymorphism at splice sites of human introns

For each of the 170,015 introns located within coding regions, we analyzed patterns of polymorphism in the vicinity of its donor splice site (last 20 bp from the upstream exon, and first 30 bp of the intron) and of its acceptor splice site (last 30 bp of the intron, and first 20 bp from the downstream exon), using polymorphism data from the 1000 Genomes Project (phase 3; ftp://ftp.1000genomes.ebi.ac.uk/vol1/ftp/release/20130502/) (The 1000 Genomes Project Consortium 2012). Derived allele frequencies were measured on the entire sample (2504 individuals from 26 populations), for all SNPs for which information on ancestral state was available. In total, our data set includes 447,659 SNPs (0.026 SNP per bp), among which 437,080 (97.6%) with DAF information.

## DATA ACCESS

Illumina read sequences generated in this study have been submitted to the European Nucleotide Archive (ENA) (https://www.ebi.ac.uk/ena) under accession number PRJEB15532. Processed data (human, paramecia) are available at http://doi.org/10.5281/zenodo.321639.

## ACKNOWLEDGEMENTS

We thank Linda Sperling for helpful comments and for sharing her analyses of the distribution of splice signals within paramecium CDSs. We thank Olivier Arnaiz for his precious help in analyzing RNAseq data. We thank Ulrich Braunschweig for kindly providing data on intron retention rates and expression levels of human genes. This work was supported by the Agence Nationale de la Recherche (ANR-12-BSV6-0017-04 INFERNO), and by the France Génomique national infrastructure, funded as part of the “ Investissements d’Avenir” program managed by the ANR (ANR-10-INBS-09). It received support under the program “Investissements d’Avenir” launched by the French government and implemented by the ANR with the references ANR-10-LABX-54 MEMOLIFE and ANR-11-IDEX-0001-02 PSL Research University. This work was performed using the computing facilities of the CC LBBE/PRABI.

